# Development and validation of an exome-wide SNP genotyping array for genomic prediction, GWAS and assessment of introgressive hybridization between black and red spruces, and transferability to white and Norway spruces

**DOI:** 10.1101/2025.11.14.688480

**Authors:** Sébastien Gérardi, France Gagnon, Nathalie Pavy, Jérôme Laroche, Simon Nadeau, Brian Boyle, André Soro, Shona Millican, Iain Thompson, Ashley Thomson, Martin Perron, Simon Bockstette, Jean Beaulieu, Patrick Lenz, Jean Bousquet

## Abstract

Introgressive hybridization plays a major role in shaping the evolutionary dynamics and adaptive potential of forest trees. In this study, we developed and validated an exome-wide bispecific SNP genotyping array (Pmr25k) for the closely related species black spruce (*Picea mariana*) and red spruce (*Picea rubens*), two ecologically and economically important North American conifers that form a widespread hybrid zone in eastern Canada. Exome capture and sequencing of pooled red spruce samples yielded over 25,000 high-quality SNPs, which were used in conjunction with a previously developed black spruce gene SNP resource of over 97,000 high-quality SNPs, to construct the bispecific genotyping array. The final array comprised 21,573 successfully manufactured SNPs, representing 14,200 distinct gene loci, of which 85% were segregating when both species were considered together. More than 4000 segregating SNPs could also be successfully used and genotyped in each of white spruce (*Picea glauca*) and Norway spruce (*Picea abies*), highlighting the conserved nature of DNA attachment sites and presence of homologous SNPs for many gene loci. The Pmr25k array thus provides an efficient and reliable high-throughput genotyping tool to investigate introgression, genetic adaptation at the gene level, and to assist genomic-based prediction for breeding and conservation efforts in boreal spruces.

## Introduction

Introgression refers to the incorporation of genetic material from one taxon into the gene pool of another through hybridization, followed by repeated backcrossing. In long-lived taxa with large effective population sizes such as perennial forest trees, introgression may persist over long evolutionary timescales and leave durable signatures in the genome (Jaramillo-Correa et al. 2009, Godbout et al. 2012). Introgression in forest trees has diverse evolutionary consequences, mainly by generating novel genetic variation and shaping adaptation. For instance, empirical evidence suggests that interspecific gene flow through introgression contributed to adaptive evolution through the transfer of new adaptive alleles and traits across species boundaries in both Angiosperms (e.g. *Quercus*: Leroy et al. 2020, Liang et al. 2025; *Populus*: Suarez-Gonzalez et al. 2018, Rendón-Anaya et al. 2021), and conifers (e.g. *Pinus*: Zhao et al. 2014, Menon et al. 2021; *Picea*: Jaramillo-Correa & Bousquet 2005, Hamilton et al. 2015, Liu et al. 2024). From a conservation perspective, introgression presents both opportunities and risks. On one hand, the incorporation of potentially advantageous alleles may provide a source of genetic variation that could enhance adaptive capacity under rapid climate change. On the other hand, introgressive gene flow from widespread taxa can erode the genomic integrity of narrowly distributed species, leading to homogenization or even extinction through genetic swamping and maladaptation (Lepais et al. 2009, Pfennig et al. 2016, An et al. 2017). The balance between adaptive potential and genetic vulnerability underscores the importance of studying introgression in forest trees, particularly within species complexes that include threatened taxa. A clearer understanding of its mechanisms and evolutionary consequences appears essential for both fundamental research and applied fields such as tree breeding and conservation genetics (de Lafontaine et al. 2015, Hamilton & Miller 2015), and even more so in the context of climate change.

Despite its importance, introgression in forest trees remains underexplored compared to model plants or agricultural species. This is particularly true for conifers, which possess some of the largest and most complex genomes among plants (De La Torre et al. 2014, Lo et al. 2024). Yet, North American boreal conifers offer an excellent model for studying introgression, owing to their wide geographic distributions and their cyclical range shifts associated with Pleistocene glacial oscillations (Hewitt 2000, Jaramillo-Correa et al. 2009), which resulted in recurrent interspecific contact. Moreover, advances in high-throughput sequencing and population genomic methods are now providing unprecedented opportunities to dissect the extent and functional consequences of introgression (Bousquet et al. 2021).

Black spruce (*Picea mariana* [Mill.] BSP) and red spruce (*Picea rubens* Sarg.) are two North American conifer species, which bear significant ecological and economic value. Black spruce is a dominant species of the boreal forest (Beaulieu et al. 2004, Prunier et al. 2011, 2012), whose broad transcontinental distribution covers most of Canada and some areas of northeastern United States (Jaramillo-Correa et al. 2004, Gérardi et al. 2010), with among the highest genetic diversity at the SNP level of genes among conifers (Pavy et al. 2025). Red spruce has a more meridional and fragmented distribution spanning from high-elevation habitats in the southern Appalachian Mountains, to more septentrional regions of northeastern North America and southern Quebec (Blum 1990).

The two species are closely related, being sister taxa in phylogenetic analyses (Bouillé et al. 2011, Lockwood et al. 2013). They hybridize naturally within their area of sympatry, which spans southern Quebec, the Maritimes, and the bordering states east of the Great Lakes in the United States (Blum 1990, Perron & Bousquet 1997). Historically, natural introgressive hybridization between black spruce and red spruce has been extensively investigated using a wide range of genetic markers. Early genetic studies based on Random Amplified Polymorphic DNA (RAPD) (Perron et al. 1995, Perron & Bousquet 1997), Sequence-Tagged nuclear DNA Site (STS) (Perron et al. 2000), and mitochondrial DNA (Jaramillo-Correa & Bousquet 2005) markers showed that introgression was prevalent in the area of sympatry and that gene flow was asymmetrical, primarily occurring from black spruce to red spruce. Perron et al. (2000) also provided evidence that black spruce and red spruce should be considered as a progenitor-derivative species pair that likely diverged during the Pleistocene, a conclusion later supported by mtDNA polymorphisms (Jaramillo-Correa & Bousquet 2003). The advent of high-throughput sequencing and genotyping technologies made SNP markers widely accessible, allowing for more refined interpretations of earlier findings. Based on hundreds of SNPs, de Lafontaine et al. (2015) showed that certain genomic regions remained impermeable to gene flow between the two hybridizing taxa, thereby maintaining species boundaries despite ongoing introgression. Using the same SNP dataset, de Lafontaine & Bousquet (2017) confirmed that gene flow occurred predominantly from black spruce to red spruce, while backcrossing was more frequent from red spruce to hybrids.

Due to their high ecological and economic importance, both species are the focus of extensive breeding programs in Eastern Canada (Mullin et al. 2011), where genomic approaches are also increasingly being integrated to support tree improvement and forest management (Lenz et al. 2017, Bousquet et al. 2021). Despite substantial progress, several key questions remain unresolved, many of which being of critical importance to forest managers, including whether introgression is spatially uniform across the sympatric area, whether exogenous selection effectively maintains species boundaries, or how introgression might be harnessed to improve tree growth, wood quality, and resilience to environmental change (Reed-Métayer et al. 2025). In this work, we describe the development of a new genotyping tool specifically designed to address these questions. We leveraged the high genomic relatedness between black spruce and red spruce (Perron et al. 2000) to develop a high-density Infinium iSelect gene SNP array (here called Pmr25k) comprising 25,000 beads and designed to genotype trees from both species simultaneously. To this end, we first performed exome capture on allopatric red spruce natural populations and generated a large catalog of red spruce gene SNPs. Using this new catalog and the black spruce gene SNP catalog previously published (Pavy et al. 2016), we primarily selected SNPs shared between the two species with the restriction of maximizing exome coverage within the limits of this SNP array. We then complemented this selection with species-specific SNPs identified in allopatric populations of each species. The genotyping success rate and preliminary results support the view that this approach appears broadly applicable to similar complexes of closely-related species for a variety of genomic monitoring and prediction applications.

## Materials and methods

### Development of a red spruce SNP resource

The step by step workflow that lead to the development of the red spruce gene SNP resource is presented in Figure 1. Twigs were collected for three allopatric populations from the southernmost parts of the species range, that showed very limited historical introgression with black spruce (de Lafontaine et al. 2015) (Table 1). Each population included 9 to 15 trees, for which DNA was extracted from needles using the DNeasy 96 Plant Kit of Qiagen (Mississauga, ON, Canada). A 550 ng DNA pool was prepared for each population, thus for a total of three genetically distinct DNA pools. All trees belonging to the same population were represented at equimolar concentrations in each pool. Libraries were synthesised with NebNext Ultra II DNA library preparation kit for Illumina (NEB) according to the manufacturer’s instructions. A final amplification step was conducted with Kapa HiFi hotstart ReadyMix 2x (Kapa Biosystems).

**Figure 1.**
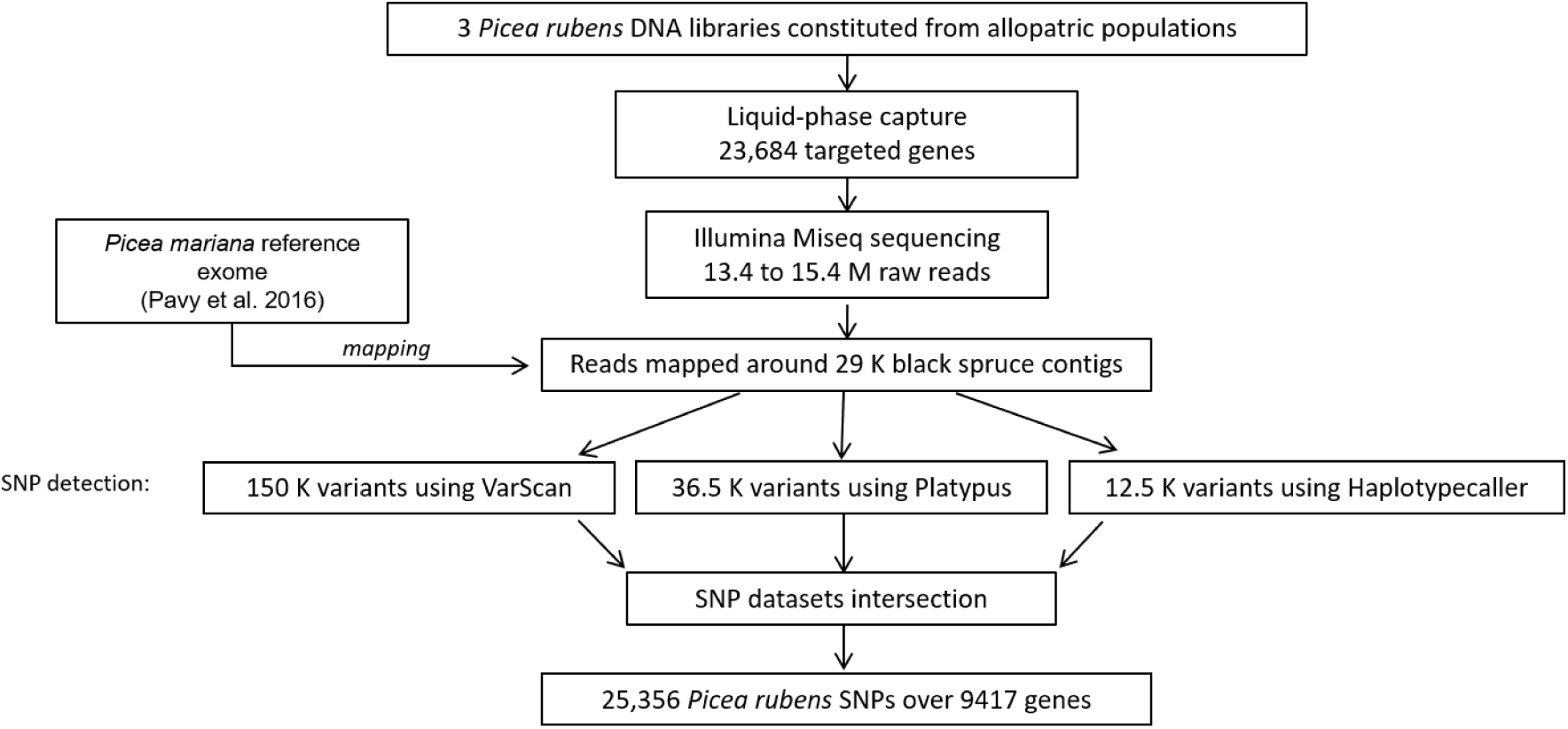
Pipeline describing the general workflow implemented to generate the red spruce SNP resource. Sequential steps consisted in performing an exome capture and sequencing, followed by mapping red spruce reads on a black spruce exome previously published (Pavy et al. 2016) and detecting SNPs using three different SNP callers. SNPs retained at the final stage had to satisfy several in-house quality filtering criteria (see Methods for more details).

**Table 1.**
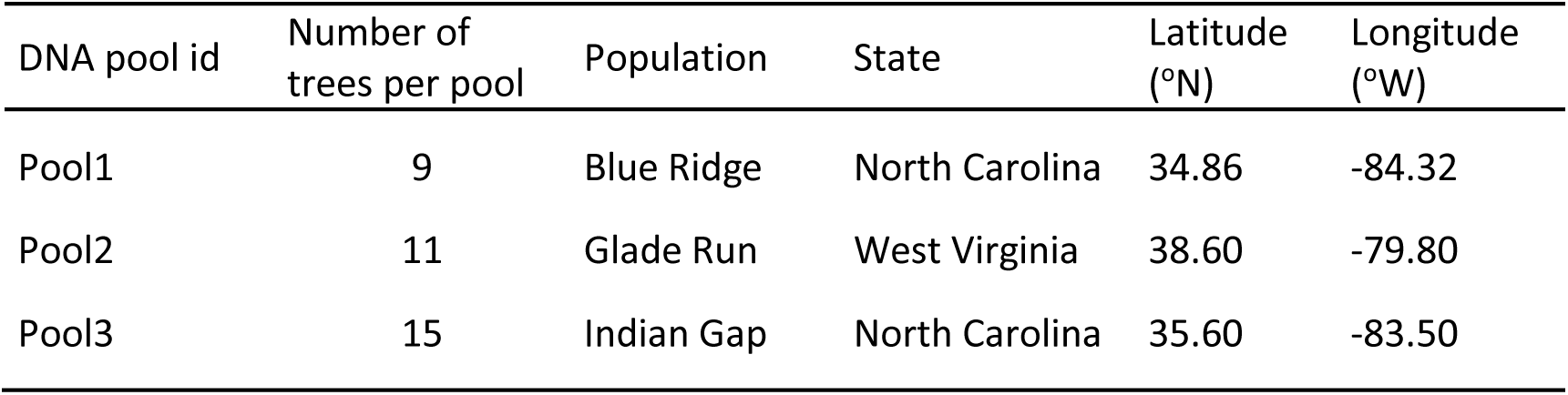
Geographic origins of allopatric red spruce DNA pool samples used in the exome capture and sequencing.

| DNA pool id | Number of trees per pool | Population | State | Latitude (°N) | Longitude (°W) |
| --- | --- | --- | --- | --- | --- |
| Pool1 | 9 | Blue Ridge | North Carolina | 34.86 | -84.32 |
| Pool2 | 11 | Glade Run | West Virginia | 38.60 | -79.80 |
| Pool3 | 15 | Indian Gap | North Carolina | 35.60 | -83.50 |

As previously described for black spruce (Pavy et al. 2016) and Norway spruce (Azaiez et al. 2018), a liquid-phase capture (SeqCap EZ developer, IRN 6089042357, OID35086; Roche Nimblegen) targeting coding region sequences of 23,864 genes was performed, using a set of probes developed on white spruce by Stival Sena et al. (2014). Following the capture step, individual libraries were sequenced at the Institute for Systems and Integrative Biology (Univ. Laval, Québec, Canada) using the Illumina MiSeq paired-end technology, which generated 300 bp reads.

Read adapters were trimmed using Trimmomatic 0.39 and sequences shorter than 80 bp were discarded. Given the close phylogenetic relatedness between black spruce and red spruce (Perron et al. 2000), red spruce reads were then aligned against a black spruce reference exome already available (Pavy et al. 2016) using BWA-MEM (Burrows-Wheeler Alignment) with a minimum seed length of 33 bp, a mismatch penalty of 10 and a gap open penalty of 100. The mapping of red spruce reads on a black spruce reference exome allowed us to establish an accurate correspondence between SNPs detected in both species.

SNP detection was carried out using three SNP callers, namely VarScan 2.3.9, Haplotypecaller 4.0.11, and Platypus 0.8.1.1. Details regarding the exact parameters used for each software are provided in Supplementary Method S1. Briefly, only biallelic non-singleton SNPs were retained and quality filtering applied post SNP calling followed the recommendations provided in Jasper et al. (2022). Particular attention was given to DP (depth) and GQ (genotype quality) filters, as these parameters had the greatest impact on genotyping success. Both were individually optimized for each SNP caller to minimize false positives (see Supplementary Methods S1). Only SNPs meeting these quality criteria and identified by two SNP callers or more were considered valid, which was also a procedure recommended by Jasper et al. (2022). We also retained red spruce SNPs identified by a single SNP caller if the SNP was previously detected in black spruce (see Pavy et al. 2016 for details regarding the production of the black spruce SNP resource).

### Development of a bispecific genotyping array

An Illumina Infinium iSelect SNP array of 25,000 beads (named Pmr25k) was developed by selecting a subset of type II SNPs necessitating one bead/SNP, and type I SNPs necessitating two beads/SNPs, from the red spruce gene SNP catalog (developed herein) and the previously published black spruce gene SNP catalog (Pavy et al. 2016). SNP selection aimed at maximizing exome coverage and therefore, no more than three SNPs per gene were retained. To assign SNPs/contigs to genes, we used Exonerate v2.4.0 to align black spruce contig sequences with homologous white spruce gene sequences (identity ≥ 95%) obtained from GCAT v3.3 (Rigault et al. 2011), the most recent and comprehensive white spruce gene catalog. SNPs located on gene contigs assigned to more than one gene, as well as those with a percent identity below 0.95, were still considered without being assigned to a single gene.

Three consecutive steps of SNP selection were conducted. We first selected ∼5000 black spruce SNPs that were successfully genotyped on previous arrays (de Lafontaine et al. 2015, Pavy et al. 2016). Some of them were also successfully genotyped on a limited subset of red spruce individuals, thus reinforcing their relevance for the development of the current bispecific array. Each of them represented a single gene and all were type II SNPs. The additional SNPs (∼18,000) were newly selected from the previously published black spruce SNP resource (Pavy et al. 2016) and the new red spruce gene SNP resource developed herein. Selected SNPs had to satisfy the following criteria: biallelic; Illumina functionality score ≥ 0.60 without neighboring polymorphisms 50 pb upstream or downstream; type II SNPs favored over type I SNPs because only one bead needed per type II SNP; minor allele frequency (MAF) ≥ 0.10; shared SNPs between black spruce and red spruce favored. Finally, we expanded exome coverage by including ∼450 interspecific SNPs located in unrepresented genes, that is, markers which were fixed for an allele in a given species while this allele had a frequency lower than 0.10 in the other species. The rationale for including these markers was that, although these sites were monomorphic or nearly monomorphic in one or the other species in their allopatric populations, they were expected to segregate in introgressed trees.

As expected, given the lower genetic diversity of red spruce (Perron et al. 2000, Jaramillo-Correa and Bousquet 2003, de Lafontaine et al. 2015, Capblancq et al. 2020), the number of genes represented in the red spruce SNP resource (Table S1) was comparatively limited relative to that of black spruce (Pavy et al. 2016), which resulted in an array design unbalanced in favor of black spruce SNPs. The final SNP list included 23,490 SNPs (including some type I SNPs necessitating 2 beads per SNP), of which 20,894 SNPs were detected in black spruce (89.0% of the array) and 8528 in red spruce (36.3% of the array), while 6376 SNPs were shared between both species.

These SNPs were distributed over 14,915 distinct gene loci. The list was submitted to the Genome Quebec Innovation Centre (Montreal, Canada) genomic platform for manufacturing the Illumina iSelect genotyping array, which was carried out by Illumina Inc. (San Diego, California, USA), according to previous excellent genotyping success rates from Illumina genotyping arrays that we have designed for a variety of spruce taxa over the past 15 years (Pavy et al. 2008, 2012, 2013, 2016, Pelgas et al. 2011, Lenz et al. 2020a, 2020b, Tumas et al. 2024).

Genotype calling was performed using the Genome Studio 2.0.5 software (Illumina Inc.). SNPs were visually inspected and genotype clusters were manually curated using the editing clusters function of the Genome Studio software with the ‘No-call threshold’ adjusted at 0.10. Two criteria were considered for calling successful SNPs: 1) segregating for more than 0.1% of individuals, and 2) SNP call rate ≥ 80%.

### Plant material and DNA extraction of trees genotyped on the SNP array

To this day, a total of 12,805 black spruce and red spruce trees were genotyped sucessfully on this new Pmr25k SNP array. These various subsets of red and black spruce genotyped trees support a wide range of objectives, from fundamental research on the geographic and genomic characterization of introgression patterns, to applied goals including the use of GWAS and genomic prediction to enhance tree breeding programs and genetic conservation efforts. For each species, the sampling was based on a number of allopatric and sympatric natural populations from Canada and the United States (total of 1384 trees), breeding populations from provincial tree improvement programs in the provinces of Ontario, Quebec, New Brunswick and Nova Scotia in Canada, as well as individuals from a 20-year-old progeny test located in the Quebec province and resulting from controlled crosses (Reed-Métayer et al. 2025) (total representing 11,421 additional trees). SNP occurrence in allopatric areas, assumed to reflect ancestral polymorphism within each species, was assessed using subsets of representative control trees sampled from these regions. For this purpose, allopatric populations of each species were defined in the strict sense as those located at least 500 km from the reported putative area of sympatry (Figure 2), to minimize the likelihood of interspecific genetic contact and natural introgression that could occur well beyond the recognized area of overlapping natural distributions. In addition to red spruce and black spruce trees, we included a limited number of trees from white spruce (*Picea glauca*) and Norway spruce (*Picea abies*) from various populations, in order to assess the rate of SNP transferability (or sharing) among congeneric taxa. Therefore, a total of 26 white spruce trees (from Pelgas et al. 2011, Beaulieu et al. 2014) and 28 Norway spruce trees (from Mottet et al. 2015) were also genotyped, representing various geographical origins for each species.

**Figure 2.**
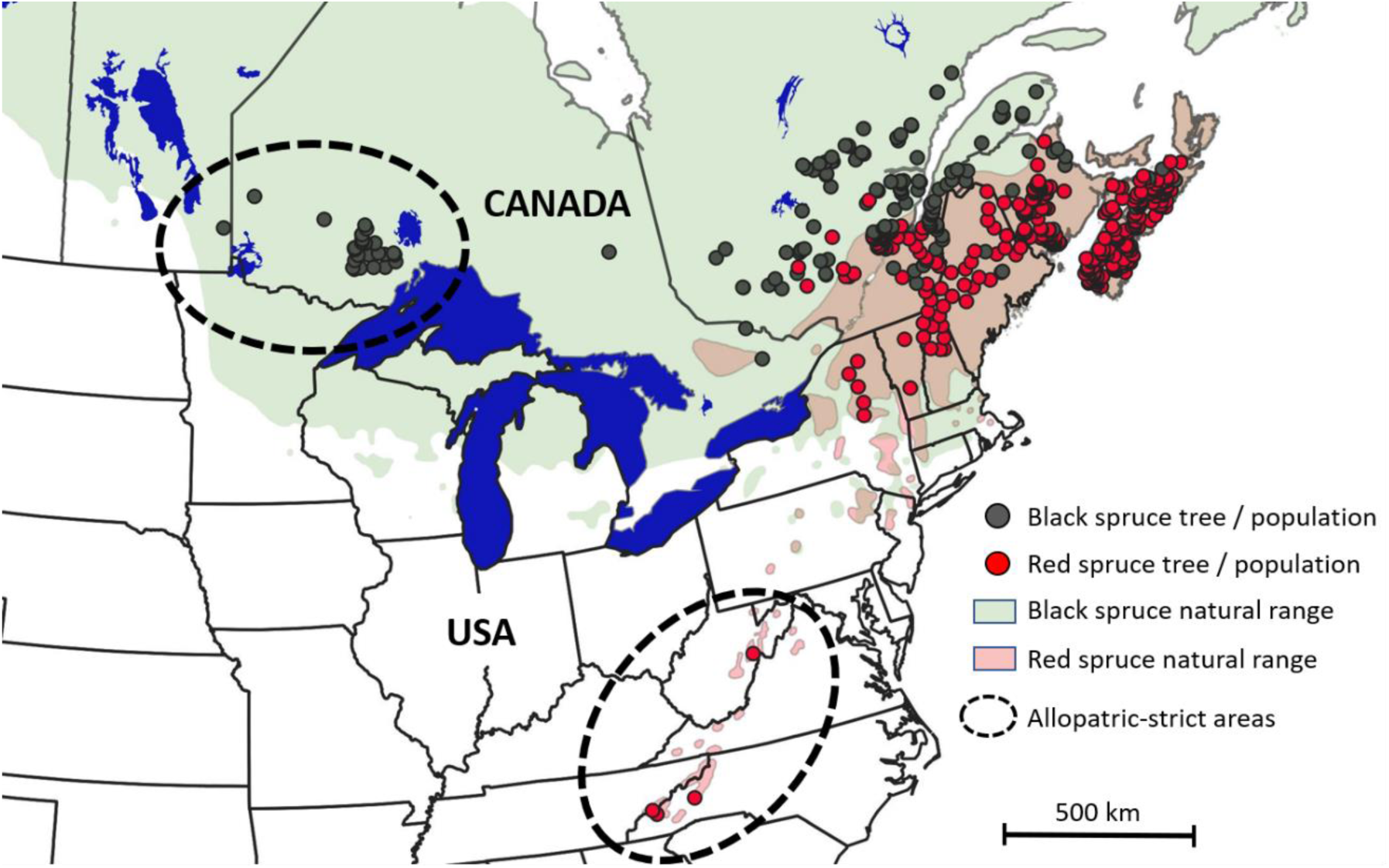
Sampling locations of 12,805 genotyped trees from black spruce (black circles) and red spruce (red circles) relative to their respective natural ranges (light green for *P. mariana* and light pink for *P. rubens*) including those from natural populations and those from breeding tests. Dashed black ellipses indicate natural populations from allopatric areas defined in the strict sense, that is, located at least 500 km away from the putative sympatric area between the two species, in order to better identify species-specific SNPs and avoid the zone of potential genetic contact and introgressive natural hybridization between species (see Methods).

In addition, the intra-assay reproducibility rate was evaluated by including two positive controls on each genotyping plate of 96 samples and comparing their genotypes obtained. These controls were identical to those used in previous spruce projects given that they were parents of controlled crosses between black spruce and red spruce (Pavy et al. 2008, Pavy et al. 2016), which allowed us to evaluate the inter-assay reproducibility for the ∼5000 black spruce SNPs reused from these previous SNP arrays.

For all genotyped trees, DNA was extracted from buds and needles using the same kit as described earlier (see *Exome capture and sequencing in the red spruce* section) and a minimum of 80 ng of DNA was used for genotyping.

### Genomic introgression index

The genomic introgression index, or ancestry coefficient, was estimated for all black spruce and red spruce trees sampled using the Bayesian algorithm fastSTRUCTURE 1.0 (Raj et al. 2014). The K value was set to 2, as the focus was primarily on species assignation rather than possible intraspecific population structure patterns. Individuals were then sorted according to their ancestry coefficients and plotted to assess genomic admixture across the sampling.

## Results and Discussion

### Red spruce exome capture and sequencing and production of a SNP resource

The MiSeq paired-end sequencing generated between 13.4 M and 15.4 M raw reads for each of the three red spruce pools (Table 2). For each pool, high-quality mapping on the black spruce exome sequence was achieved for approximately 30% of the total number of red spruce reads with mapped reads representing ∼29,000 black spruce gene contigs (Table 2).

**Table 2.** Red spruce exome capture sequencing and mapping metrics onto the black spruce exome.

| DNA pool id | Raw reads | Mapped contigs | Mapped reads | % mapped reads |
| --- | --- | --- | --- | --- |
| Pool1 | 13.4 M | 28,992 | 3,805,375 | 28.6 |
| Pool2 | 13.8 M | 28,995 | 3,817,498 | 28.5 |
| Pool3 | 15.4 M | 29,008 | 4,317,213 | 28.4 |

Following the SNP calling procedure and the subsequent quality filtering steps, VarScan detected ∼150,000 high-quality variants, whereas ∼36,500 and ∼12,500 were detected by Platypus and Haplotypecaller, respectively (Figure 1). In their study, Jasper et al. (2022) reported that using overlapping variant lists among callers is an effective strategy to minimize false positives. After applying this approach to our SNP datasets, we obtained over 9 K red spruce SNPs identified by more than one SNP caller. We also identified ∼19,000 red spruce SNPs identified by a single SNP caller, but shared with the black spruce SNP resource previously developed by Pavy et al. (2016). Hence, the final new red spruce SNP resource included a total of 25,356 distinct SNPs (Supplementary Table S1) matching 9417 black spruce gene contigs and 8053 white spruce gene homologs.

The total number of gene SNPs identified herein among red spruce pools (∼25.5 K) appeared rather limited compared to what was previously obtained for black spruce with similar criteria of high-quality gene SNPs (total of over 97,000 SNPs; Pavy et al. 2016). While this result likely reflects that red spruce is less diverse than black spruce, it may have been exacerbated by the comparatively lower sequencing effort for red spruce in the framework of our experimental goals, which primarily focused on the production of a bispecific genotyping array, rather than maximizing the detection of rare SNPs in red spruce. For instance, we used fewer samples in the DNA pools sequenced at the exome capture step because we intentionally limited our tree selection to the most allopatric red spruce provenances so to avoid as much as possible introgression. Such a procedure would limit the discovery of low MAF alleles in natural red spruce populations. Moreover, the short-read MiSeq sequencing obtained herein was not complemented by a sequencing technology yielding longer reads, as was the case for black spruce where 454 GS-FLX-Titanium sequencing was used in conjunction with Illumina HiSeq sequencing (Pavy et al. 2016). Finally, red spruce sequencing reads were mapped on the black spruce exome sequence (Pavy et al. 2016), which was critical to perform an efficient SNP selection for the final goal of constructing a bispecific SNP array, but it was not optimal for maximizing polymorphism detection in red spruce since it likely led to the exclusion of many informative reads. Thus, these reported contrasting levels of detection of exome SNP abundance between red spruce and black spruce should not be used as a direct and unbiased measure of their comparative exome SNP diversity, although the trend towards reduced SNP diversity in red spruce is certainly real.

### Genotyping array reproducibility, success rate, and distribution of SNP minor allele frequencies

The SNP array manufacturing process resulted in 21,573 correctly manufactured SNPs representative of 14,200 distinct gene loci, including about 1298 type I SNPs necessitating two beads per SNP and the rest, 20,275 type II SNPs necessitating only one bead per SNP. Illumina probe synthesis failed for 8.2% (1917) of the total number of SNPs initially supplied to the genotyping platform, a rate of failure within expectations according to previously manufactured Illumina Infinium iSelect SNP arrays for several conifers (Howe et al. 2013; Pavy et al. 2013, 2016, Lenz et al. 2020a, 2020b, Tumas et al. 2024). Positive controls replicated across the 152 genotyped DNA plates enabled the estimation of intra-assay genotyping reproducibility at 99.99% for the Pmr25k array. Inter-assay reproducibility based on ∼5000 SNPs recovered from the PmGP1 array (Pavy et al. 2016) reached 98.3%, confirming the high reliability of the genotyping results obtained herein and reproducibility of the technology.

A total of 18,272 segregating SNPs met the quality criteria used to determine the genotyping success rate (see *Development of a bispecific genotyping array* section for additional details) (Supplementary Table S2), representing an overall genotyping success rate of 85% (Table 3) across all black spruce and red spruce trees genotyped on the Pmr25k array. These segregating SNPs represented 12,438 distinct gene loci based on the exome catalog of black spruce SNPs (Pavy et al. 2016), of which 11,572 genes were homologous to white spruce genes from the GCAT 3.3 gene catalog (Rigault et al. 2011). The most challenging aspect of developing the Pmr25k array was to identify as many shared SNPs between black spruce and red spruce as possible so that the genotyping array could be used efficiently for various purposes in both species. Given that the capture probes were designed using white spruce genes (Stival Sena et al. 2014), it is likely that most successful targeted gene sites were conserved and shared by ancestry among species (see Bouillé & Bousquet 2005), as they would have predated the speciation of the three spruce taxa, thereby contributing to the high genotyping success rate observed in both species.

**Table 3.** SNP segregating rate for black spruce and red spruce tree groups genotyped on the Pmr25k SNP array, considering only trees with call rate ≥ 0.85.

| Group | Number of trees | Number of segregating SNPs | Percent of total segregating SNPs<sup>1</sup> | Number of distinct gene loci |
| --- | --- | --- | --- | --- |
| Black spruce trees from allopatric-strict natural populations (see Figure 2) | 32 | 16,658 | 91.2 | 11,698 |
| Red spruce trees from allopatric-strict natural populations (see Figure 2) | 33 | 8463 | 46.3 | 6298 |
| Black spruce trees from natural populations from potential contact zone | 364 | 18,171 | 99.4 | 12,388 |
| Red spruce trees from natural populations from potential contact zone | 955 | 18,230 | 99.8 | 12,423 |
| Black spruce breeding populations from Ontario and Québec | 5804 | 18,125 | 99.2 | 12,363 |
| Red spruce breeding populations from New Brunswick and Nova Scotia | 4633 | 18,160 | 99.4 | 12,381 |
| Black spruce x red spruce hybrid common garden | 984 | 18,028 | 98.7 | 12,295 |
| Total number of black spruce and red spruce trees | 12,805 | 18,272 | 100 | 12,438 |

<sup>1</sup> In reference to the total number of 18,272 segregating SNPs of the entire Pmr25k SNP array.

Overall, only 30 out of the 12,805 trees genotyped with the new SNP array had a call rate below 85%, the pre-established threshold for excluding samples. When examining only the 32 individuals from black spruce allopatric-strict populations (Figure 2), the proportion of segregating SNPs was 91.2% (Table 3). Despite the limited sample size, this high polymorphism rate observed suggests that this subset of allopatric trees captured most of the diversity present in the species complex. On the other hand, the proportion of segregating SNPs for the 33 allopatric-strict red spruce trees (Figure 2) was approximately half, at 46.3% (Table 3). While this result suggests that allopatric red spruce trees are genetically less diverse than their black spruce counterparts with many more non-segregating SNPs, it may also reflect at least in part the fact that the array design was biased in favor of black spruce SNPs (see *Development of a bispecific genotyping array* section for more details). Hence, inferences on genetic diversity within species and differences between species drawn from this study should be interpreted with caution, though several trends point to a lesser genetic diversity detected in red spruce than in black spruce, which would correspond to the proposed progenitor-derivative speciation hypothesis between these two species, black spruce being the progenitor and red spruce, the derivative species (Perron et al. 2000, Jaramillo-Correa & Bousquet 2003).

The detailed MAF distribution of SNPs based on individuals from allopatric-strict and sympatric natural populations is presented in Figure 3. Figure 3A shows that, within allopatric populations, the number of non-segregating SNPs (MAF = 0) was substantial when each species was analyzed separately, but considerably reduced when both species were analyzed jointly. This pattern indicates that only a limited number of SNPs were non-segregating across both species, while many SNPs were truly species-specific in allopatric populations, thus exhibiting allelic divergence between species as these SNPs were segregating (that is, polymorphic) in one species but non-segregating (that is, monomorphic) in the other. The number of non-segregating SNPs was also especially high in red spruce allopatric populations (over 10,000 SNPs; Figure 3A) for two likely reasons: 1) sampled red spruce allopatric populations were geographically isolated at higher elevation in the southern Appalachian mountains and have been generally reported as genetically depauperate (Perron et al. 2000, Jaramillo-Correa & Bousquet 2003, Capblancq et al. 2020), and 2) a greater proportion of the SNPs included on the array were originally identified in black spruce, relative to red spruce. In sympatric populations, the number of non-segregating (monomorphic) SNPs was low in all groups, whether the species were considered individually or jointly (Figure 3B). This result suggests that gene flow and introgression actively take place between the two species in their sympatric area, leading to an increase in shared polymorphisms and number of segregating SNPs. Although less prominent that in allopatric populations, red spruce still showed a MAF distribution of SNPs skewed towards lower values, likely indicating lower effective population size and/or persistent asymmetric introgression, which agrees with previous studies focusing on this species complex (de Lafontaine & Bousquet 2017). Altogether, although red spruce SNPs exhibited lower MAFs than black spruce on average, the number of completely non-segregating SNPs on the array remained low, thereby reinforcing the utility of this genotyping tool for future fundamental and applied studies.

**Figure 3.**
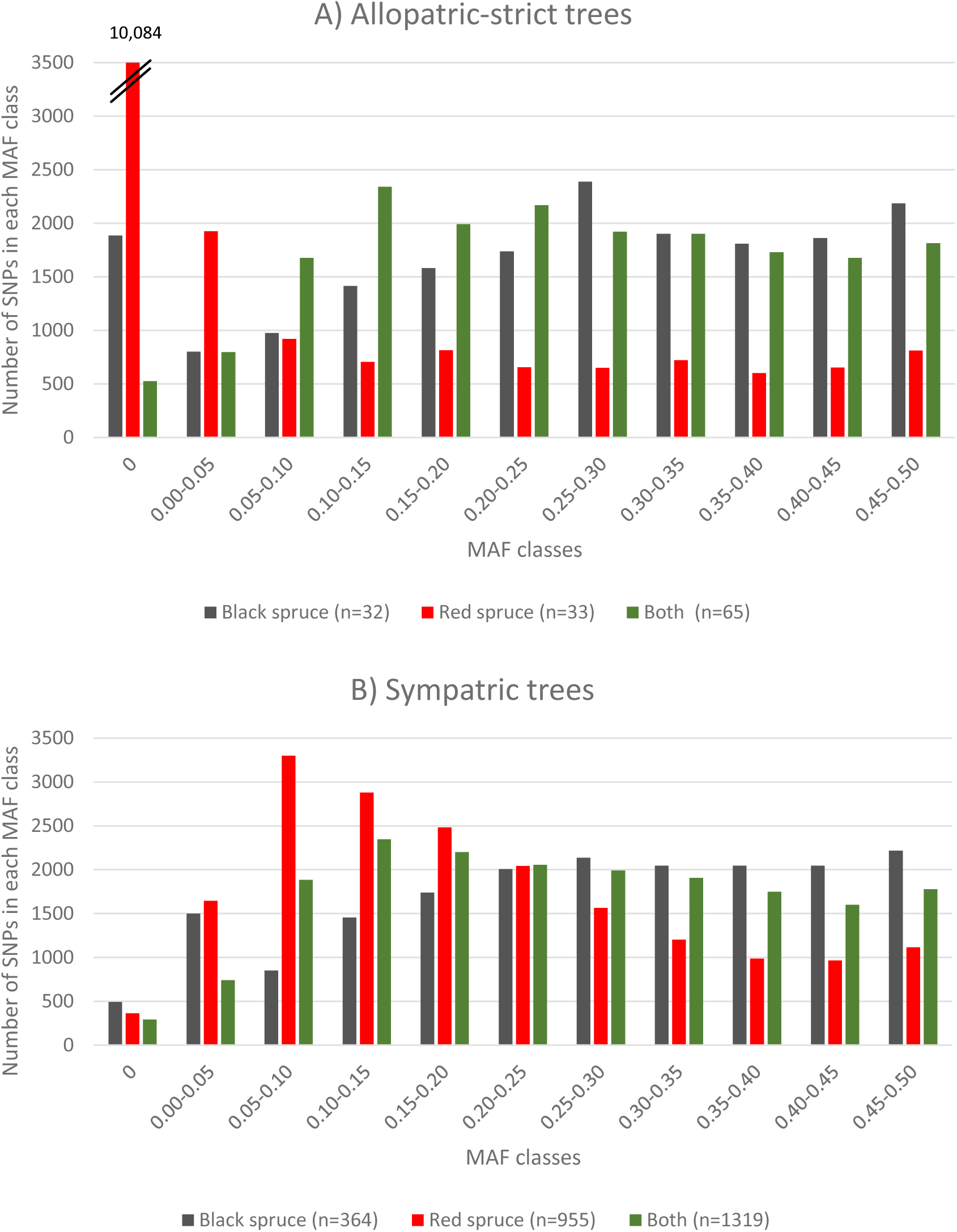
Distribution of minor allele frequencies (MAF) for successful SNPs from the Pmr25k SNP array for black spruce trees (black bar), red spruce trees (red bar) as well as for both species together (green bar); A) for trees from allopatric-strict natural populations, and B) for trees from sympatric natural populations.

### Transferability of successfully arrayed black spruce and red spruce SNPs towards other spruce species

Using this SNP array especially designed for the phylogenetically closely-related black spruce and red spruce, we tested the transferability rate towards the more phylogenetically distant and reproductively isolated white spruce and Norway spruces. The genotyping of 26 white spruce and 28 Norway spruce trees from various provenances on this SNP array resulted in 4621 segregating SNPs for white spruce (covering 3888 distinct gene loci) and 3497 segregating SNPs for Norway spruce (covering 3082 distinct gene loci). These SNPs represented 25.3% and 19.1% of the total number of segregating SNPs identified for black and red spruce, respectively. This transferability rates are higher than those observed when black spruce (17.6%) and Norway spruce (12.5%) were genotyped using previous white spruce SNP arrays (Pavy et al. 2013).

Several factors may account for this marginally increased transferability: (1) among the 5K Pmr25k SNPs carried over from previous genotyping black spruce SNP arrays (see Materials and Methods), 560 were already known to be polymorphic in white spruce; (2) our selection criteria favored SNPs shared between black and red spruce, thus increasing the proportion of ancestral variants likely shared across other spruce species; and (3) a larger number of white spruce and Norway spruce trees were tested compared to the earlier test of interspecific transferability between reproductively isolated congeneric taxa using white spruce SNP arrays (Pavy et al. 2013), hence resulting in more power for detecting SNPs with MAF values ≤ 0.10 in white spruce and Norway spruce. However, the lesser rate of transferability towards the Eurasian Norway spruce, as previously reported (Pavy et al. 2013), likely reflects its larger phylogenetic distance from the black spruce - red spruce lineage, compared to the North American white spruce (Bouillé et al. 2011).

This relatively high SNP transferability across phylogenetically distant congeneric spruce taxa suggests that a significant proportion of genetic variants are conserved across spruce species from shared common ancestry, as previously reported from the sequence analysis of homologous nuclear genes (Bouillé & Bousquet 2005). Moreover, the transferability rate reported here is likely underestimated, as the relatively small sample size of white spruce and Norway spruce trees genotyped with this new SNP array limited the detection of rare SNPs in these taxa, especially for those with MAF ≤ 0.05. Therefore, these empirical observations highlight the utility of multi-species SNP panels for genotyping congeneric species in the genus *Picea*, an observation replicating the trend observed for *Eucalyptus* sp. (Grattapaglia et al. 2015). The identification of thousands of segregating SNPs in both white spruce and Norway spruce using a large SNP array originally designed for black and red spruce opens possibilities for comparative genomic studies across spruce species and potentially broader genomic applications in the population genetics, genetic conservation or tree breeding fields of research.

### Genomic introgression index

The FastSTRUCTURE analysis revealed a continuous distribution of genomic ancestry coefficients across red and black spruce individuals (Figure 4), with a large proportion of trees showing intermediate values. This pattern of widespread genomic admixture is consistent with extensive hybridization and gene flow between the two species, in agreement with previous findings highlighting the close evolutionary relationship between these species and the role of historical and ongoing gene flow in shaping their genetic landscape (Perron and Bousquet 1997, Perron et al. 2000, Jaramillo-Correa & Bousquet 2003, Jaramillo & Bousquet 2005, de Lafontaine et al. 2015, 2017, Capblancq et al. 2020).

**Figure 4.**
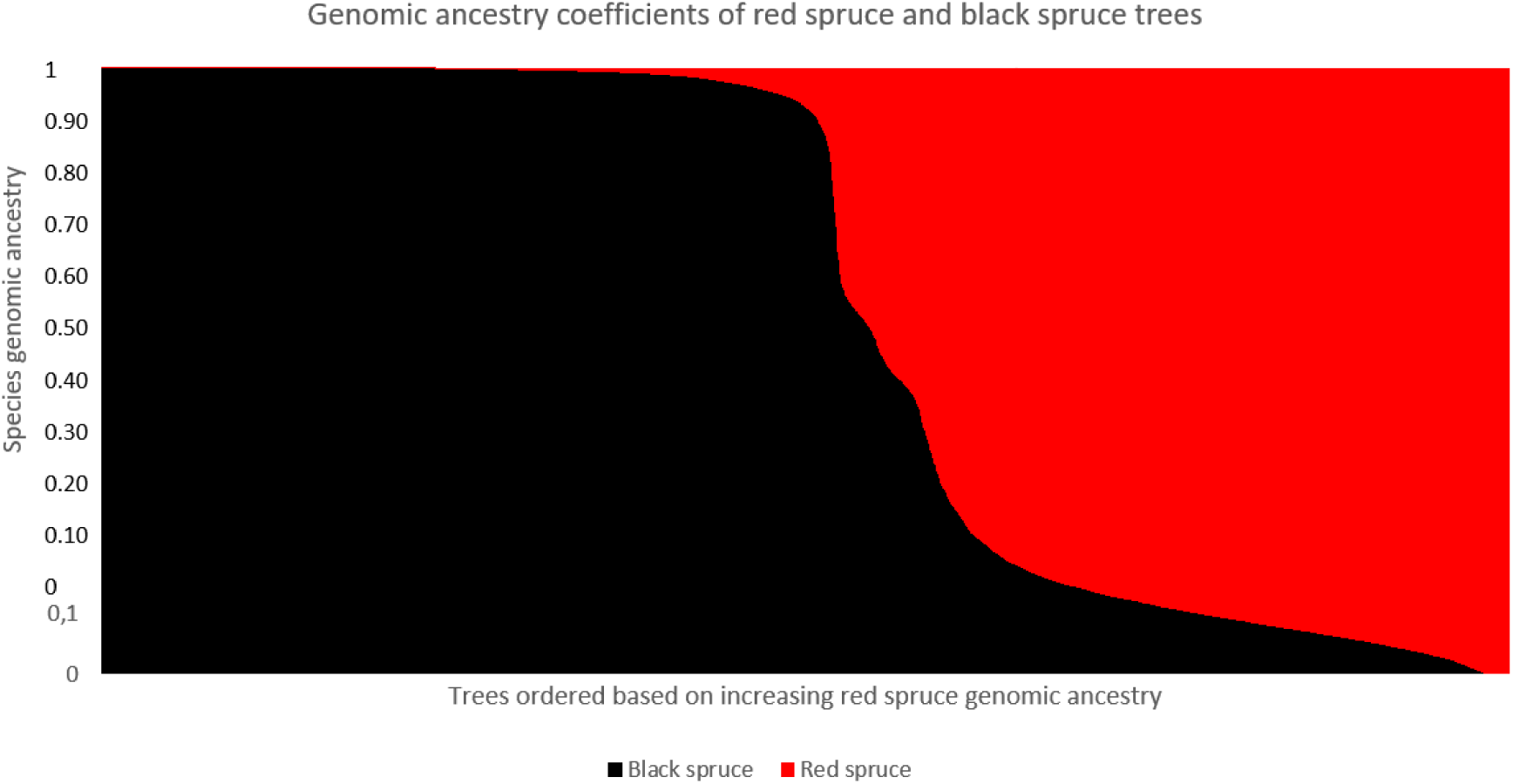
Genomic ancestry coefficients (also called genomic introgression index) estimated using FastSTRUCTURE for all black spruce and red spruce trees genotyped on the new Pmr25K SNP array. Each vertical bar represents a single tree, and the color proportions reflect the estimated proportion of that tree’s genome assigned to black spruce (black) and red spruce (red). Trees are ordered along the x-axis based on increasing red spruce ancestry.

## Conclusions

The development of the Pmr25k bispecific SNP array represents a major advance for genomic research in the *P. mariana* x *P. rubens* species complex. The SNP array achieved high genotyping success rate and reproducibility, confirming its reliability and cost-effectiveness for large-scale applications in multiple breeding programs and for other more fundamental applications.

Although allopatric red spruce exhibited lower overall SNP polymorphism than black spruce, consistent with its narrower distribution and reported reduced genetic diversity, substantial shared variation between the two species was detected. Patterns of SNP minor allele frequency and ancestry coefficients revealed extensive genomic admixture and gene flow, providing clear evidence of widespread introgression across their known sympatric area and beyond. The relatively high transferability of SNPs to other more phylogenetically remote and reproductively isolated *Picea* species further highlights the conserved nature of sequence and polymorphism at many gene loci, and the broader utility of this genomic resource across the genus. Overall, this study demonstrates that black spruce and red spruce form an interconnected species complex shaped by ongoing natural introgressive hybridization. The Pmr25k SNP array should thus provide a powerful and versatile platform to monitor gene flow in natural populations, investigate the genomic architecture of introgression, and explore its adaptive significance under changing environmental conditions. Beyond advancing our understanding of evolutionary questions, this new genomic tool should be key in supporting the development of genomic-assisted breeding strategies in boreal spruce taxa.

## Acknowledgements

Funding for this work was provided by a grant from the Genomic Application Partnership Program (GAPP) of Genome Canada and Genome Quebec to J. Bousquet and P. Lenz, leaders of the FastTRAC II spruce genomics project, and all collaborators and personnel from government and private partners who gathered the spruce samples from natural populations, provenance tests and breeding populations. Thanks to Elizabeth Bronswijk for editing a near final draft of the manuscript. We also thank Elizabeth McKinley and Larissa Robinov from the New Hampshire Department of Natural and Cultural Resources, as well as Randall Morin from the United States Department of Agriculture, for kindly providing valuable data and insights that supported the design of our sampling strategy for populations located in the United States.

## Supplementary information

### Supplementary Method S1

Description of the parameters used for each SNP caller and quality filtering criteria applied subsequently.

### Varscan 2.3.9

#### Vcf file was first generated using the following criteria

Min coverage: 2

Min reads2: 2 (remove singletons)

Min var freq: 0.0

Min avg qual: 10

P-value thresh: 0.01

#### Variants were subsequently filtered using the following criteria

Retain only biallelic SNPs polymorphic in at least one pool

DP > 10

GQ > 20

### HaplotypeCaller 4.0.11

#### Vcf file was first generated using the following criteria

genotyping-mode DISCOVERY

stand-call-conf 20

#### Variants were subsequently filtered using the following criteria

Retain only biallelic SNPs polymorphic in at least one pool

window 35 -cluster 3

Remove singletons

Hardfiltering, retain:

DP > 10

QD > 2.0

QUAL > 50

SOR < 3.0

FS < 60.0

MQ > 56.0

MQRankSum > -12.5

ReadPosRankSum > -8.0

GQ > 20

### Platypus 0.8.1.1

#### Vcf file was first generated using the following criteria

genIndels=1 --minReads=2 --maxReadLength=500 --maxVariants=3 --skipDifficultWindows=1 --minMapQual=20 --minBaseQual=20 --minGoodQualBases=20 --badReadsThreshold=10 --rmsmqThreshold=20 --hapScoreThreshold=20 --filterReadsWithUnmappedMates=0 --filterReadsWithDistantMates=0 --filterReadPairsWithSmallInserts=0 --trimAdapter=0 --maxGOF=20 --sbThreshold=0.01 --scThreshold=0.95 --hapScoreThreshold=15 --filterDuplicates=0 --verbosity=2 --minFlank=10 --filterVarsByCoverage=0 --filteredReadsFrac=0.7 --minVarFreq=0.01

#### Variants were subsequently filtered using the following criteria

Retain only biallelic SNPs polymorphic in at least one pool

Remove singletons

QUAL > 20

DP > 30

**Supplementary Table S1.** Description of the red spruce SNP resource

See excel file

**Supplementary Table S2.** Detailed information on SNPs genotyped on the Infinium Pmr25k SNP array

See excel file

